# Rational Exploration of Fold Atlas for Human Solute Carrier Proteins

**DOI:** 10.1101/2021.08.21.457230

**Authors:** Tengyu Xie, Ximin Chi, Bangdong Huang, Fangfei Ye, Qiang Zhou, Jing Huang

## Abstract

Solute carrier (SLC) superfamily is the largest group responsible for transmembrane transport of substances in human cells. It includes more than 400 members which are organized into 65 families according to their physiological function and sequence similarity. Different families of solute transporters can adopt the same or different folds that determine the working mechanism and reflect the evolutionary relationship between SLC members. Analysis of structural data in literatures shows that there are 13 different folds in the solute carrier superfamily covering 40 families and 343 members. To further study their working mechanism, we systematically explored the SLC superfamily to look for more folds. Based on our results, at least three new folds were found for SLC superfamily, and one was experimentally verified in the SLC44 family. Our work has laid a foundation and provided important insights for the systematic and comprehensive study on the structure and function of solute carriers.

## Introduction

Membrane proteins occupy about 30% of the human proteome, playing important roles in signaling transduction, energy generation and utilization, and mass transporting. Solute carrier (SLC) superfamily is the largest group that catalyzes the transmembrane transporting of substances in human cells^1,2^. There are more than 400 SLC members that can be divided into 65 families according to their physiological function and sequence homology. The SLC transporters typically contain several transmembrane (TM) helices that form specific folds to mediate the cross membrane transporting of the substrates. The folds of SLC members usually define their working mechanisms and imply the evolutionary relationship between them even when the sequence similarities are low. It is well known that distant-homology proteins can share the same fold, with human SLC superfamily as a typical case with high sequence diversity but relatively low structural diversity. Exploring new folds in a rational way can thus accelerate the establishment of the structural landscape, and ultimately facilitate our understanding on the functions of the SLC superfamily. In this work, we have systematically studied the folding of SLC transporters by computationally screening and then experimental verification and established a full structural atlas for the human SLC superfamily.

### Protocol to identify new folds

Identification of new protein folds in a genome has been attempted in the early 21st century, where the probability for a sequence adopting a new fold is estimated by statistical models established from sequence-based classifications^3-6^. Some secondary structure prediction results are used^7^, however, these methods rely mainly on sequence similarity.

The recent surge in the accurate prediction of protein structures, which is benefited from the efficient utilization of coevolution information and deep neural networks, provides an opportunity to rethink the task of identifying new protein folds from their sequences. Well-designed neural networks can realize the mapping of multiple sequence alignments to structures by learning from structure database. This is due to the inherent ability of the residual neural network and the attention-based network to infer pairwise relationship. The idea was originally implemented in RaptorX-contact^8^ and reached a climax in AlphaFold^9,10^ that have made a great progress recently. RoseTTAFold^11^ was developed using a similar end-to-end network architecture and achieved a comparable accuracy of structure prediction to AlphaFold. The high accuracy of AlphaFold makes it possible to offer meaningful predicted structures to biologists in human genomics^12^. Before the merge of AlphaFold, we had focused on the SLC fold atlas and attempted to provide more knowledge about the function of human SLC superfamily by integrating predicted structures from multiple different methods.

We established an approach to rationally find new folds in the human SLC superfamily (Fig. 1). Starting from all sequences of the SLC superfamily, multiple advanced methods were used to extract accurate structural information at three levels, including 3D structure coordinates, 2D contact maps, and 1D secondary structure contents. This information was then compared with known protein structures in protein data bank (PDB)^13^ using a carefully designed score to measure the fitness of a sequence compared to all known folds (detailed in the Method section). The methods were applied to investigate the structural landscape of the human SLC superfamily.

**Fig. 1.**
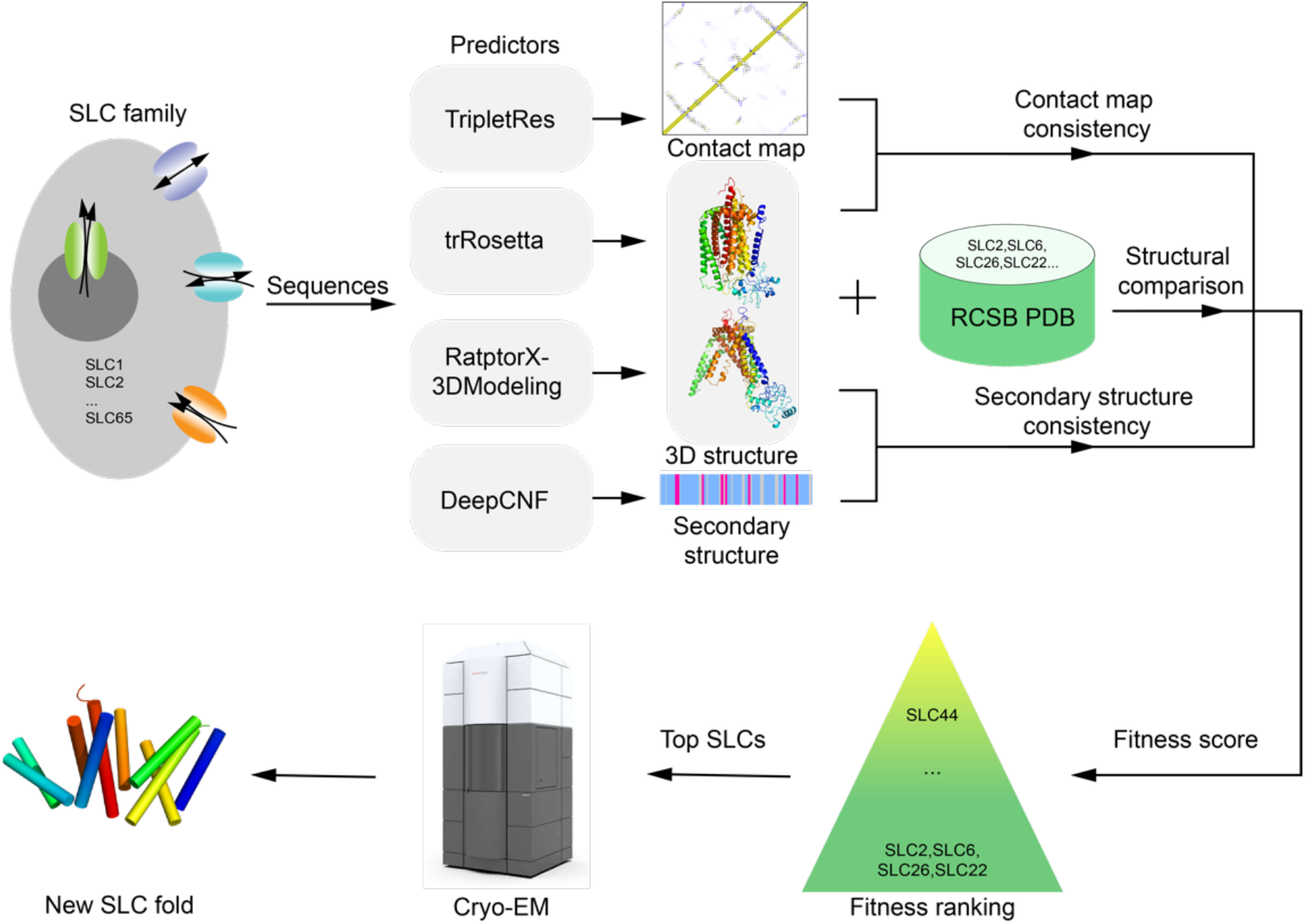
The discovery strategy of new fold of human SLC superfamily.

To ensure that the fitness score can distinguish new folds based on predicted information, analysis was performed using the knowledge of the SLC superfamily. We checked and confirmed that the SLC members with high scores have had their own or homologous structures (Table S1 and Fig. S1B). We also validated the calculation protocol by assuming that only SLC members with LeuT fold are included in experimentally known structure database while other SLC members have no experimental structures. Under this assumption, most of SLC members have very low scores which indicates that they are not LeuT fold (Fig. S1C).

During the process of our research, trRosetta^14^ and RaptorX-3DModeling^15^ were adopted as 3D structure predictors in our protocol with which a new fold SLC44 was discovered. After that, AlphaFold and RoseTTAFold were released, which can be adopted as better predictors to look for other new folds of SLC superfamily at the next iteration of our protocol. Furthermore, to better establish the SLC fold atlas, we referred to the prediction results from AlphaFold and RoseTTAFold for a few SLC members, for which the predicted structures generated by previous methods were not good enough.

### SLC fold atlas

Literature and database searching indicate that 13 folds are identified for 40 families covering 343 members in human SLC superfamily or their homologs among the PDB entries till February 2021^16-18^, while for the remaining 25 families covering 81 members, the folds are not determined (Table S2, Fig. 2B). To obtain a more completed overview of the human SLC family, we analyzed their folds by combining our prediction results with the experimental structural information. Using DaliLitev.5^19^, 424 human SLC members were classified into 29 clusters. We survey the common characteristics of each cluster and manually confirm most of clusters as reasonable folds, while some neighboring clusters with the same fold were merged. Among 424 human SLC members, 415 SLC members with more than 5 TMs were classified into 23 sorts of folds, while the other remaining 9 SLC members with no more than 5 TMs were classified into 4 classes according to the numbers of TM helices (Fig. 2A and Table S2). For each specific fold, the representative experimental structure from PDB database or predicted structure generated by AlphaFold are shown in Movie S1. We note that the members belonging to the same family adopt the same fold, indicating that the folds are consistent with the sequence-based families but offer more general function-related information.

**Fig. 2.**
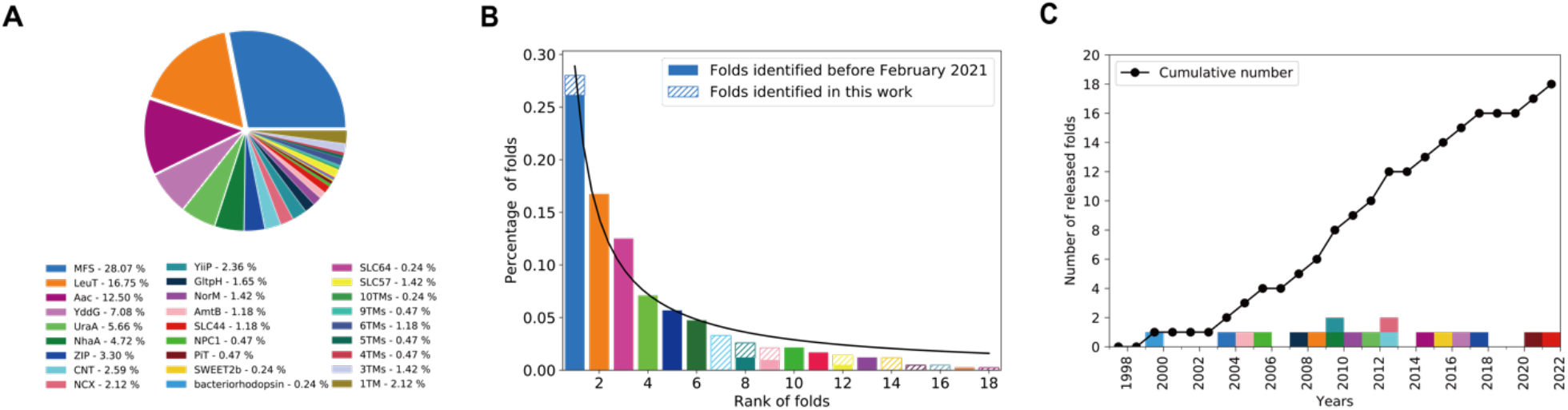
The overview of SLC folds. (**A**) The percentages of confirmed folds in SLC superfamily. (**B**) The percentages of folds before February 2021 (solid color block) and after our calculation (shaded color block). The percentages follow the Zipf’s law. (**C**) The number of released folds in each year (color blocks) and the cumulative number of released folds along years (black dot line). The released year for each fold is the earliest year of all PDBs belonging to the fold. The color of bars in b or blocks in c corresponds to the color legend of (**A**).

Based on known folds till February 2021, our calculation identified 10 more folds. In addition, some previously unclassified SLC members were classified into three existing folds (Fig. 2B). For example, four SLC families (SLC52, SLC59, SLC60, and SLC61) are newly recognized as MFS (major facilitator superfamily) (Table S2). SLC8 and SLC24, phylogenetically belonging to *δ* family^17^, were structurally classified to NCX in our work, which indicates that the structure-based classification is compatible with the phylogeny-based classification.

We further investigated the fold distribution of the human SLC members. As shown in Fig. 2B, the relationship between the percentages of the folds and corresponding ranking follows the Zipf’s law, similar to the power law observed in genome^20^. Almost half of SLC proteins (44.81%) belongs to the MFS and LeuT folds. Besides, the released years of the 18 folds are shown in Fig. 2C. New folds were discovered at a rate of about one per year from 2003 to 2012 and at a slowed down rate in recent years, indicating the fold atlas is nearly complete.

### Characteristics of SLC folds

Among the 23 human SLC folds, 21 folds are composed of two or more repeating units that contain 2-7 TM helices in a repeat and are internally correlated with each other via an inverted or rotated pseudo-symmetrical axis, which is a common feature for the SLC folds (Fig. 3, Fig. 6). There are 6 folds containing 5 TM helices and 5 folds containing 3 TM helices in their repeating unit, which are the most common number of TM helices in a repeating unit at fold level. Among the 13 folds with inverted repeats, 8 folds are composed of a core and a scaffold domain. The folds with rotated repeats usually have no core and scaffold domains. There are some folds (GltpH, CNT, and PiT) that contain a pair of half-transmembrane hairpin in their core domain. Some SLC members consist of multiple (usually double) duplicates of the fold of the other members in their corresponding family. For example, SLC14A2 contains two duplicates of the AmtB fold linked by a long linker, while AmtB fold is adopted by the other members in SLC14 (Fig. 4A). In addition, some SLC members exert their function in homo-oligomers, the folds of which consist of multiple chains such as SLC31, a trimer, forming a fold with 9 TM helices (Fig. 4B). The combination of different folds in a SLC member is also observed. The fold of SLC30A5 is composed of the YiiP and the YddG folds that are adopted by SLC30 and SLC35, respectively, as shown by the structure predicted by AlphaFold (Fig. 4C).

**Fig. 3.**
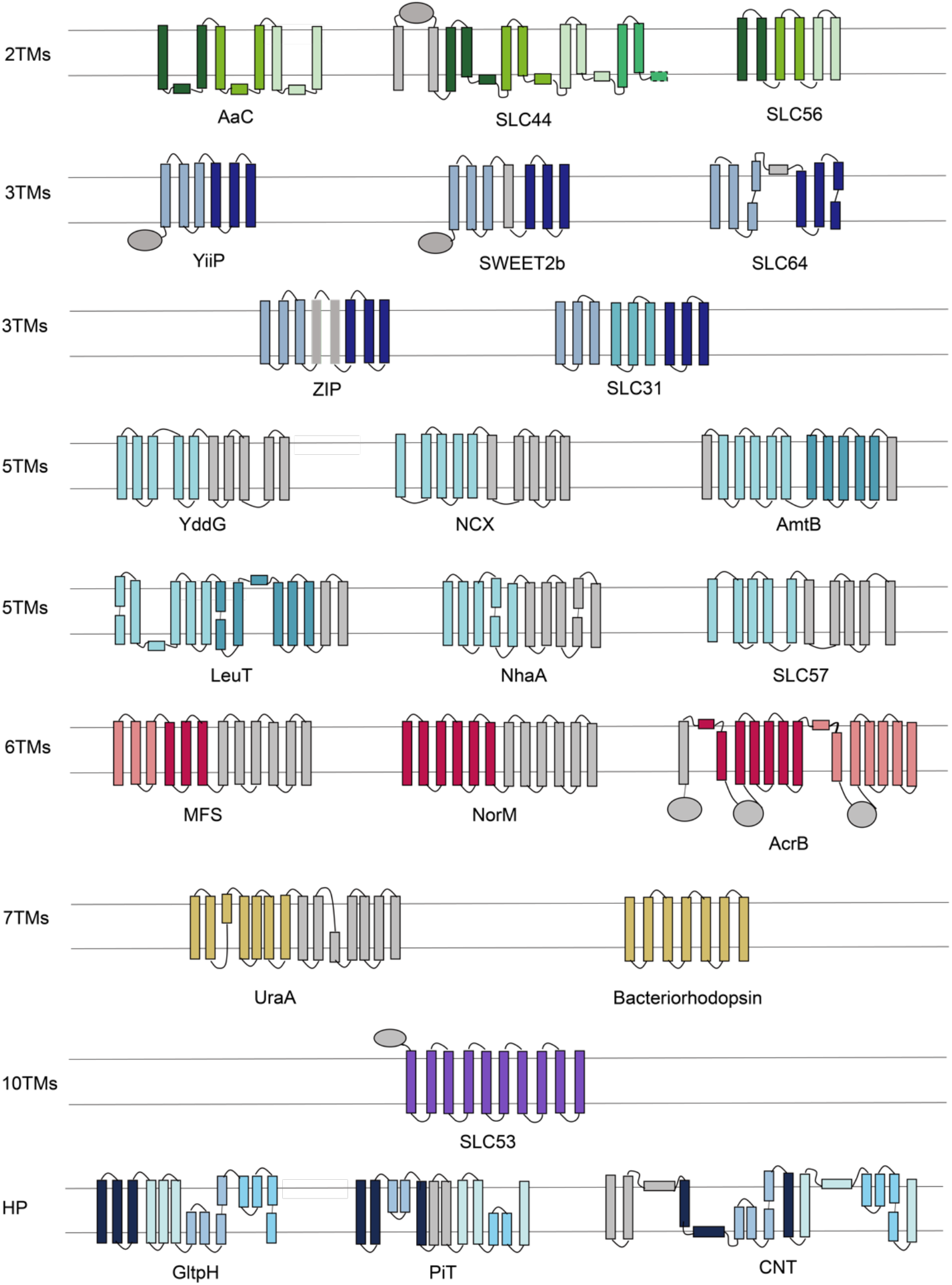
Cartoon representation of SLC folds. These folds are listed according to their number of TM helices in a structural repeat unit.

**Fig. 4.**
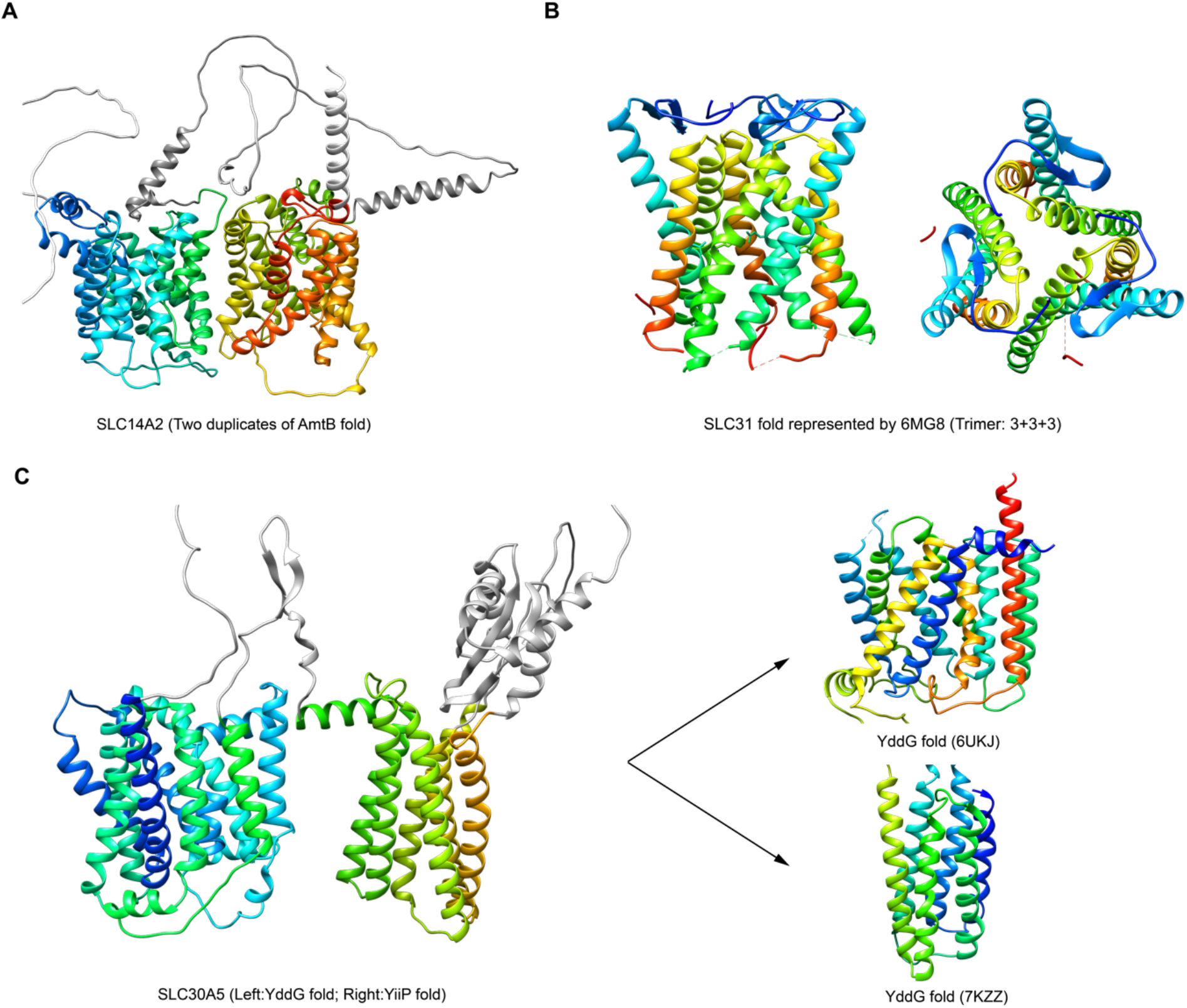
Folds of SLC14A2, SLC31, and SLC30A5. (**A**) SLC14A2 adopts the two duplicates of the AmtB fold. (**B**) The fold of SLC31 is composed of three chains, each of which has 3 TM helices. **(C)** SLC30A5 is composed of the YiiP and the YddG folds. The predicted structures of SLC14A2 and SLC30A5 are generated by AlphaFold.

SLC57 and SLC64 are new folds that have not been experimentally verified. Their predicted structures by AlphaFold are similar with those predicted by trRosetta^14^ and RaptorX-3DModeling^15^, which is convincing for further analysis. According to the ranking results, all members of SLC57 family were not in top positions and were predicted to be structurally most similar with sodium-pumping pyrophosphatase (PDB ID: 4AV6), but their sequential TM helical arrangements of the two proteins are different. SLC57 forms “5+5” inverted repeats with TM 3 inserted in the between of TM 1 and TM 2. As for SLC64 with only one member SLC64A1, which is ranked as 11^th^ (Table S3), there is no similar structures in database found. The TMs 1-3 and TMs 4-6 of SLC64A1 are folded as “3+3” inverted repeats linked by a long TM helix, which is parallel to the membrane plane. Experimental results are in demand for validation for these computational results. SLC44 is a new fold validated by experiments, which is introduced in the next section.

### SLC44 is a new fold in SLC protein superfamily

We applied our prediction protocol to 424 human SLC proteins and hypothesized that SLC members with low fitness scores might have new folds. The fitness score distribution overall is approximate to a normal distribution (Fig. S1A). The first version of calculation results indicated that the members of the SLC20 family (SLC20A1 and SLC20A2) probably adopted new folds. The later published PDB structure of a homology of SLC20 from *Thermotoga maritima* adopts a new fold, named as PiT fold^21^ (Fig. 2A). The ranking results were updated, which still indicated the members of SLC44 and SLC64 family probably adopted a new fold. We further chose SLC44A1 for structural validation using single-particle cryogenic electron microscopy (cryo-EM). SLC44A1 sequence was cloned and purified. The cryo-EM samples of SLC44A1 were prepared for data acquisition and processing (Fig. 5A, Fig. S3). Finally, a cryo-EM map of SLC44A1 was obtained at a resolution of 3.86 Å, with the resolution of TM1-2 and TM5-9 of SLC44A1 reached 3.7Å (Fig. S2B).

**Fig. 5.**
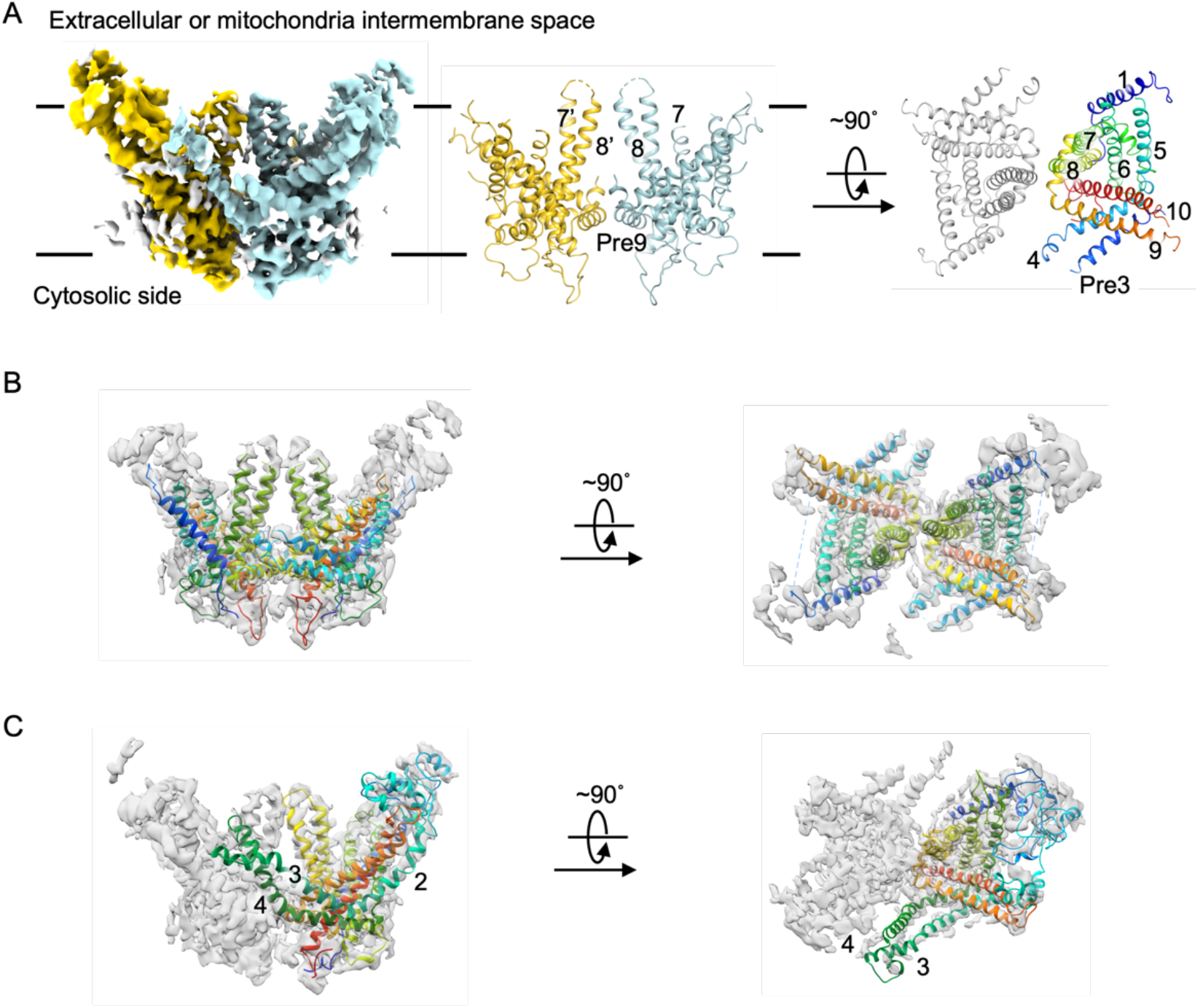
SLC44A1 adopts a new fold. (**A**) The electron density map and model of SLC44A1. The protein is captured in dimer state with TM8 and helix connecting TM8 and TM9 (Pre9) contributing the major dimerization interface. (**B**) The alignments of SLC44A1 model and its density map. The side view and top view are shown separately. (**C**) Docking result of the structure of SLC44A1 predicted by AlphaFold2 to the electron density map. The predicted structure is well-fitted to the map.

SLC44A1 has 10 TMs arranging as a corolla shape that contains 4 pairs of TM hairpins formed by TM3/4, TM5/6, TM7/8, and TM9/10, respectively. The resolution of cryo-EM map is good enough to locate most of the side chains of the transmembrane and cytosolic soluble region. The predicted structure fits the high-resolution cryo-EM map well and help to position TM2-4, where the corresponding regions in the experimental map do not have high enough resolution (Fig. 5C). The TM3/4 hairpin has different topology than the other three hairpins that assemble a rotated repeating pattern. The TM1 and TM2, which are in proximity to TM5/6 and TM9/10 hairpins in the peripheral region of TM domain, respectively, are linked by a long loop that folds as a domain. To further confirm that human SLC44A1 is a new fold, we aligned the cryo-EM structure with all structures in the PDB database by TM-align^22^. The highest TM-score is 0.46 and the corresponding protein is a sulfate permease *CysZ* ^23^ (PDB ID: 6D9Z), indicating the two structures hold different folds and SLC44A1 corresponds to a new fold.

In addition to the protein structure of SLC44A1, three regions were observed in the density map where substrates were located. The substrate in the first one was recognized as a cholesterol molecule. Its binding pocket constitutes of 12 residues from TM1, 5, 6, and 7 (Ile36, Cys39, Glu337, Trp340, Thr341, Phe398, Ile399, Cys402, Thr406, Val432, Ile436, and Leu440), which forms a hydrophobic environment to hold the cholesterol molecule. The two remaining substrates were likely to be choline or choline-derivatives, which is related to the function of SLC44A1 as a choline transporter. However, further investigation is necessary to confirm their exact chemical structures. One of the binding pockets is composed of 5 residues from TM 8-10 (Lys491, Tyr495, Asn533, Asp533, and Glu501), while the other one of 6 residues from TM5-6 and TM10 (Pro338, Thr341, Leu400, Glu403, Cys585, and Ser588).

The cryo-EM structure was also compared with predicted structures by four methods (trRosetta, RaptorX-3DModeling, AlphaFold, and RoseTTAFold) for SLC44A1. Overall, the four predicted structures hold the same fold with the cryo-EM structure (TMscores>0.5), which shows high consistency (Fig. S4). Although the two methods trRosetta (TMscore=0.67) and RaptorX-3DModeling (TMscore=0.76), adopted in our protocol, performed worse than AlphaFold (TMscore=0.95) and RoseTTAFold (TMscore=0.80) on SLC44A1, our fitness score still can offer reasonable ranking results and find new folds successfully (Fig. S4), demonstrating the robustness of the approach. The predicted structure by AlphaFold is almost the same as the cryo-EM structure while the one by RoseTTAFold seems to be in another state, where TM2 and TM7-10 are much closer with each other. Transporters undergo several conformational states to function such that the alternative states could be functionally important and will be further investigated. These results demonstrate that the developed computational approach can be used to identify proteins with new folds and thus broaden our knowledge on the sequence-structure-function relationship of proteins.

## Discussion

Proteins are the major component in cells that fulfills the life activities. Classification of the SLC superfamily is usually performed based on the sequence homology, while the sequence-structure-function relationship is central to understand the working mechanism of proteins. We present a full description on the structural atlas of human SLC proteins in this work, which will enhance our understanding in membrane transport. Our results indicate that it is possible to rationally discover new protein folds by combining computational and experimental information, allowing to systematically study the sequence-structure-function relationship for a whole protein superfamily. The similar methods can be applied to other membrane superfamilies and other species.

The SLC members usually work in single chain that folds as single domain in TM regions. There are several exceptional cases. SLC54 and SLC55 transport substrates by forming multiple chains and SLC30A5 has two TM domains. Prediction of single domain or chain of SLC55 and SLC30A5 cannot give complete insights of their functions. The information of the numbers of chains in those complexes is necessary and robust 3D structure prediction methods for multiple domains and multiple chains would be helpful ^11^.

In addition, structure clustering reveals relationship among all folds of the human SLC superfamily (Fig. 6). The major features for each fold were labeled, such as inclusion of the inverted or rotated repeats and whether it contains the half-transmembrane hairpin or not. It is worthy to note that the folds with similar features cluster closely. The 3TMs repeat unit in MFS also exists in its neighboring ZIP, YddG, and SLC57 folds. All of these four folds are different in the number of TM helices and the assembly manner of the repeating units. The SLC members of the AaC, YiiP, SLC44, and SLC56 folds with rotated repeats are located adjacently. So do the LeuT, NhaA, GltpH, NCX, UraA, and SLC57 folds that are composed of core and scaffold domains. The relationship among the SLC folds revealed based on the structural similarity may facilitate the mechanism study of the human SLC superfamily.

**Fig. 6.**
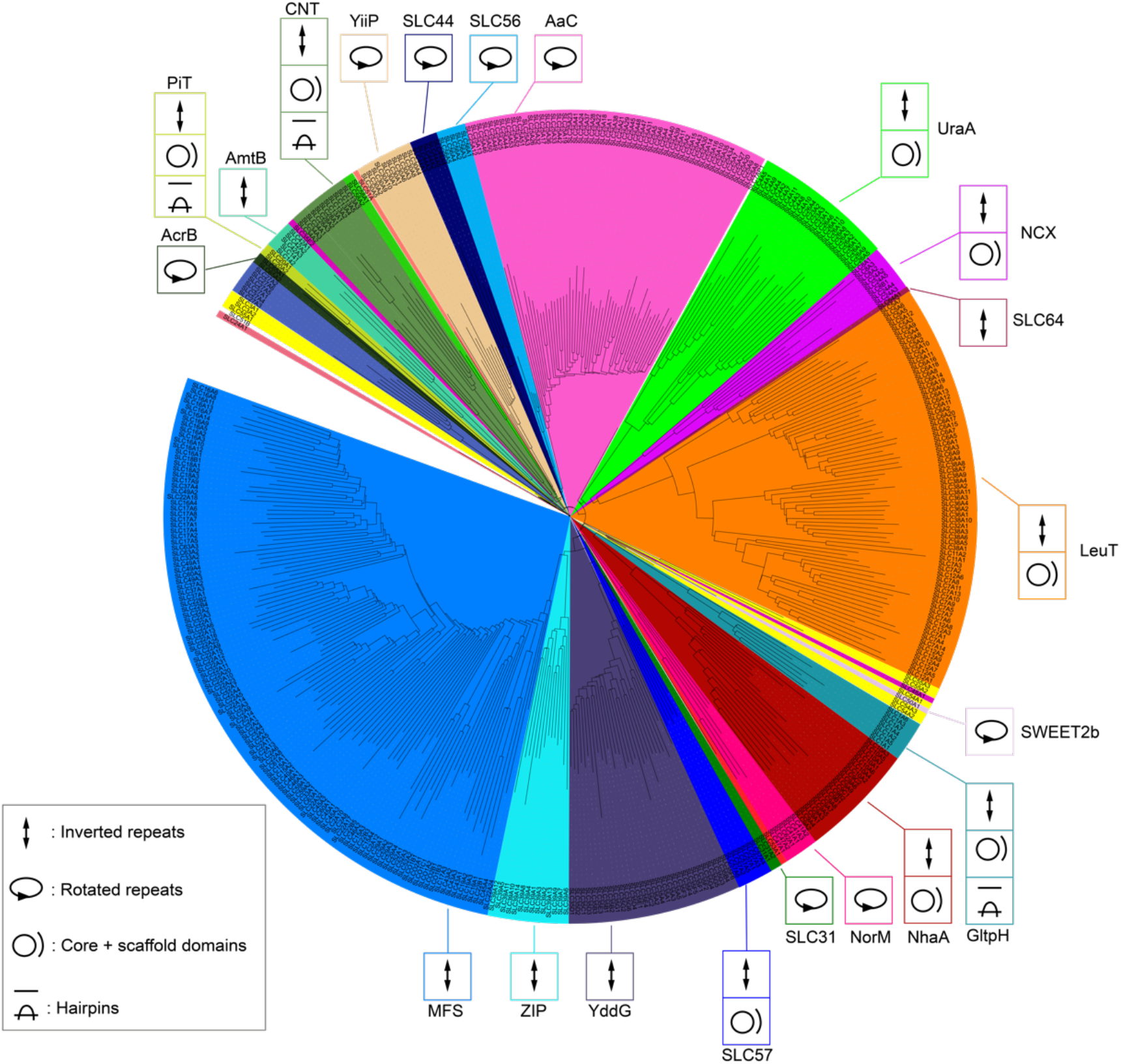
Classification result of SLC structures. The SLC members are colored according to the corresponding folds.

## Materials and Methods

### Datasets of SLC sequences and experimental structures

There are 65 SLC gene families with 458 different human transporter genes, including repeated genes with different isoforms (such as SLC14A1_UT-B1 and SLC14A1_UT-B2). As 27 SLC genes have no available amino acid sequences and 6 SLC members are isoforms, these 33 SLC members were excluded in our calculation. In total, 425 SLC genes was included in structure prediction phase. *SLC7A5P1*, which is a pseudogene, is not considered in fold recognition. Thus, 424 SLC genes were included as the dataset of SLC sequences. As the purpose is to recognize the overall fold pattern, only TM regions are subjected to structure prediction. Sequences that are 20 residues away from TM-regions are deleted during prediction. The TM regions are recognized by TOPCONS^24^.

To identify new folds, structural comparison between SLC members to all known folds is achieved by comparing SLC members to all available structures in the database. In PDB database, many proteins have multiple experimental structures, which are redundant for the structural comparison. Thus, we made use of sequence clustering data (https://cdn.rcsb.org/resources/sequence/clusters/bc-100.out) to get only one structure for a unique sequence. This reduces the 560,635 protein chains in the PDB database as of February 2021 to 97,033 sequences that are retained after clustering. We further reduced computational cost by filtering out protein sequences with no-TM regions, which resulted in 4,812 structures to be used in the computational protocol for new fold identification.

### Fitness score

To find SLC members with new fold, structural information is predicted at three levels that are 3D structures, contact maps, and secondary structures. 3D coordinate structure is the most informative and directly used to compared with known protein structures. Contact map and secondary structure prediction are less informative but more accurate to compute, so they are used to account for possible errors in 3D structure prediction. The information at each level can be predicted by customized and multiple methods. The three levels of information are integrated to a fitness score *S*, which reflects the dissimilarity of a target sequence to all experimentally determined structures and is defined as follow:

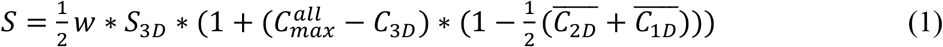

where *w* is a weight factor, *S*_3*D*_ is the similarity of the predicted 3D structures to all known structures (Eq. 2), *C*_3*D*_ is the maximum similarity of *M* 3D prediction methods for one SLC (Eq. 3),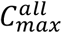 is the maximum similarity value of *C*_3*D*_ among all SLC members, 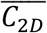 is the averaged consistency of predicted 2D contact maps to all predicted 3D structures (Eq. 4 and 5), 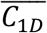 is the averaged consistency of predicted 1D secondary structures to all predicted 3D structures (Eq. 6). The fitness score *S* is a number ranged from 0 to 1 and a higher value means a protein sequence is more like known structures.

One of advantages of the fitness score is that it can integrate different prediction methods. Once new and more advanced computational methods are newly released, the fitness score can directly make use of them and keep pace with the development of predicted methods. In this work, we used two structure prediction methods: trRosetta^14^ and RaptorX-3DModeling^25^. TripletRes^26^ and DeepCNF^27^ are adopted to predict contact map and secondary structure, respectively. Predicted 3D structures are compared with all known protein structures in the PDB database through TM-align^22^. TM-align rely on structural alignment instead of sequence alignment and a TM-score value greater than 0.5 indicates that the two compared structures have similar folds^22^. Due to the inevitable inaccuracy in predicted 3D structures, two strategies are adopted to compensate the errors. One is more than one 3D protein structure prediction methods are adopted, resulting multiple predicted 3D structures for one SLC. These predicted structures are compared to all known structures, and the corresponding maximum TM-score value *S*_3*D*_ reflects the similarity of the predicted 3D structures to all known structures. The similarity *S*_3*D*_ is defined as follow:

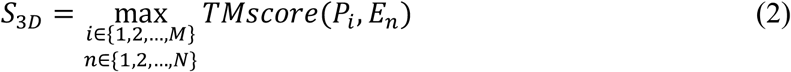

where *M* is the number of 3D structure prediction methods adopted, *N* is the number of experimental structures in the structure database. Assume *M* 3D structure prediction methods are adopted, the predicted structure *P*_*i*_ of *i*th (*i* ∈ {1,2, …, *M*}) method for one target sequence is compared to *N* experimental structures (*E*_*n*_, *n* ∈ {1,2, …, *N*}) in the database by TM-align.

In addition, the consistency of these predicted structures is also key information. If these predicted structures are similar with each other, the structures will be more reliable, and the comparison result *S*_3*D*_ based on predicted 3D structures is more convincing. Thus, the maximum similarity of all predicted 3D structure pairs (*C*_3*D*_), representing the consistency of *M* prediction methods, is integrated to *S*, where the similarity *C*_*ij*_ of a predicted structure pair *P*_*i*_ and *P*_*j*_ (*i, j* ∈ {1,2, …, *M*}) are calculated by TM-score. The definition of *C*_3*D*_ is as follow:

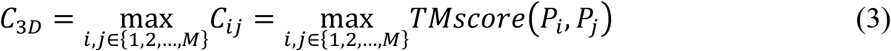

As secondary structures and contact maps can be predicted in general with higher precision, the two types of predicted information are also included in the inference of new folds. A good, predicted 3D structure should be consistent with the predicted secondary structure and contact map. Thus, 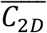 reflects the consistency of *M* 3D predicted structures to predicted contact maps and is integrated to the fitness score. Its definition is as follows:

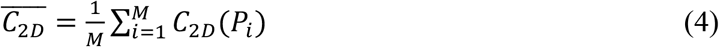

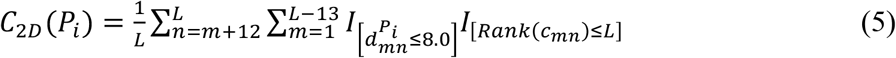

where *L* is the sequence length of a target protein, *m* and *n* are residue indices, *c*_*mn*_ is the contact probability predicted by an advanced method for the residue pair (*m* and *n*), 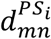 is the Euclidean distances of the residue pair in the predicted structure *P*_*i*_, the function *I*_[*condition*]_ is equal to 1 when *condition* is satisfied otherwise 0. For one predicted structure *P*_*i*_, only top *L* medium- and long-contacts (|*m* − *n*| ≥ 12) are checked. If the Euclidean distances 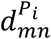 of their C_β_ atoms are no more than 8.0 angstrom and the predicted contact probabilities *c*_*mn*_ are ranked in top *L* (*Rank*(*ν*_*mn*_) ≤ *L*), the contacts of the residue pairs are consistent. So, a larger 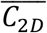 indicates the consistency of contact map is more consistent. *c*_*mn*_ is obtained from TripletRes in this work.

Similarly, the consistency of *M* predicted structures to predicted secondary structures is integrated to the fitness score. The hit rate of the secondary structures of predicted 3D structures compared to predicted secondary structures is defined as follows:

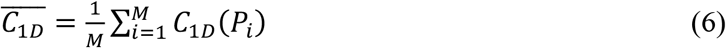

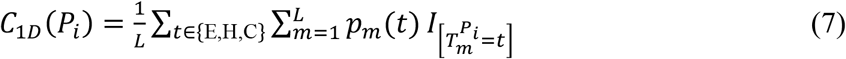

where E, H, C are types of secondary structures, *t* is one of the three types, *m* is the residue index, *p*_*m*_(*t*) is the possibility of the *m*th residue to be the secondary type *t*, 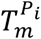 is the secondary structure type of *m*th residue of the predicted structure *P*_*i*_. DSSP^28^ is adopted to obtain the secondary structure of 3D structure *P*_*i*_. Predicted secondary structure is obtained by DeepCNF. The consistency of secondary structures is higher if 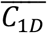 is close to 1.

Finally, the last term *w* will reweight the fitness score when the 3D structural consistency is good (*C*_3*D*_ ≥ 0.5). Its weighting down or up is determined by the similarity of predicted structures to the structural database *S*_3*D*_. Its definition is as follow:

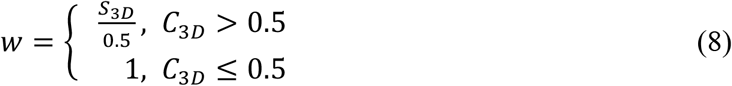

### Analysis on the fitness score

Several analyses were performed to evaluate the rationale of the fitness score *S*. All 425 members in SLC family are evaluated using the fitness score. The 15 SLC members with the highest fitness scores (Table S1) have their own or homologous structures, which are recognized by finding the most similar structures to the predicted 3D structures from all known structures. The SLC members with experimental structures are logically known folds and should have high fitness scores. Among all SLC members, 28 SLC genes have experimental structures containing transmembrane regions. Compared with all SLC members, the scores of these 28 SLC members are higher than 0.4 (Fig. S1A-B). That means these proteins are less likely to have new folds and illustrates the fitness score can recognize known folds.

Next, we assume that all known structures are SLC members with the LeuT^29^ fold, rather than all structures in PDB database and recalculate the scores for all SLC members. In principle, all other folds like MFS are different from LeuT and should be recognized as new folds with low fitness score values. As shown in Fig. S1C, most of scores of SLC members are less than 0.2 while only small percentage of SLC members have high score values, which confirms that the score can distinguish new fold and known fold. Finally, an interesting observation is that SLC20 was listed in the top position when the ranking result was firstly obtained in July 2020. Later, the experimental structure 6L85 of SLC20 was released in September 2020^21^, which confirmed it to be a new fold. All these validate the ranking protocol in this work.

### SLC members cluster determination

We use 3D structure alignment-based DaliLite.v5^19^ to identify SLC fold patterns. The structural similarity of each pair of SLC members is calculated by the intramolecular *Cα* − *Cα* differences in a set of optimized structurally equivalent residue pairs from predicted structures.

### Expression and purification of human SLC44A1

The full length sequence of human *SLC44A1* was amplified by PCR from human cDNA library, and subcloned into pCAG vector with C-terminal His and Flag tandem tags. The plasmid was purified devoid of endotoxin before transfecting HEK293F cell lines for protein overexpression. Approximately 1.5 mg plasmids were incubated with 3 mg polyethylenimines (PEIs, Polysciences) in 50 mL SMM 293T-II medium (Sino Biological Inc.) for 30 min before transfection. Cells were cultured in Multitron-Pro shaker (Infors, 130 rpm) under 37°C for 48hrs before collection and resuspension in Lysis Buffer (20 mM Hepes, pH 7.4 and 150 mM NaCl). Then the crude extraction was applied with 1.5% DDM, 0.3% CHS and 0.1875% soybean lipid (Anatrace) in 4°C overnight after addition of 1.3 mg/ml aprotinin, 5 mg/ml leupeptin, 1 mg/ml pepstatin (Amresco), and 0.2 mM PMSF (Sigma). After centrifugation at ∼25000 g for 1 hour, the supernatant containing solubilized SLC44A1 protein was load to anti-FLAG M2 affinity gel (Sigma), then washed by Lysis Buffer with 0.02%DDM and 0.004% CHS. The target protein was eluted by 0.2 mg/ml FLAG peptide and load to Ni-NTA affinity gel (Qiagen). After washed by Lysis Buffer supplied with 0.02% GDN and 0.004% CHS, the protein was finally eluted by 230 mM imidazole and further purified through size exclusion chromatography (SEC, Superose 6, 10/300, GE Healthcare) with Lysis Buffer containing 0.001% LMNG, 0.002% GDN and 0.0004% CHS. The peak fractions were checked by SDS-PAGE analysis and concentrated to ∼10 mg/ml for cryo-EM sample preparation.

### Cryo-EM sample preparation, data acquisition and procession

To prepare cryo-EM samples, aliquots (3 μL) of the concentrated protein were placed on glow-discharged holey carbon grids (Quantifoil Au R1.2/1.3) and blotted for 5 s, then flash-frozen in liquid ethane cooled by liquid nitrogen with Vitrobot (Mark IV, Thermo Fisher Scientific). 100 mM choline chloride and 0.1% FOM (Anatrace) was added before cryo-EM sample preparation. Data was collected in 300kV Titan Krios equipped with Gatan K3 detector and GIF Quantum energy filter. Movie stacks were automatically collected using AutoEMation^30^ at a defocus range from -2.2 μm to -1.2 μm in super-resolution mode at a nominal magnification of 81,000×. Each stack was exposed for 2.56 s with an exposure time of 0.08 s per frame, resulting in a total of 32 frames per stack. The total dose rate was approximately 50 e^-^/Å^2^ for each stack. The stacks were motion corrected with MotionCor2^31^ and binned 2-fold, resulting in a pixel size of 1.087 Å/pixel. Meanwhile, dose weighting^32^ was performed. The defocus values were estimated with Gctf^33^.

Particles were automatically picked using Relion 3 ^34^. After 2D classification, particles were subjected to global angular searching 3D classification against an initial model generated from similar protein by Relion 3. Then, these particles were selected by multi-reference 3D classification. After 3D auto-refinement with the map from good classes^35^, the dataset was subjected to multi-reference local 3D classification Higher regularization parameter was applied to contain more high-resolution information in 3D classification. The selected particles were further optimized by 3D non-uniform refinement in cryoSPARC. The resolution was estimated with the gold-standard Fourier shell correlation 0.143 criterion^36,37^ with high-resolution noise substitution^38^. Refer to Fig. S2-3 for details of the data collection and processing.

### Model building of human SLC44A1

The initial model of SLC44A1 was built by MDFF^39^, with the predicted structure from RaptorX-3DModeling as starting point and was guided by a previously obtained lower resolution density map. With improved resolution at 3.86 Å, we used PHENIX^40^ and COOT^41^ to modified the model. Each residue was checked manually in COOT. Several segments of the sequence were not modeled because of the invisibility of the corresponding density in the map. The dimer structure model was built by C2 symmetry applied through PHENIX and real-space-refined, then adjusted manually in COOT. Structure refinement was performed with PHENIX with secondary structure and geometry restraints to prevent structure overfitting. To monitor the overfitting of the model, the model was refined against one of the two independent half maps from the gold-standard 3D refinement approach.

## Acknowledgements

We thank the cryo-EM facility and the high-performance computing center of Westlake University for providing supports. This work was supported by the National Key R&D Program of China (2020YFA0509300) from Ministry of Science and Technology of China, the National Natural Science Foundation of China (projects 31971123, 32022037, 21803057, 32171247), and Zhejiang Provincial Natural Science Foundation of China (LR19B030001).

## Author contributions

J.H. and Q.Z. conceived this work. T.X., X.C., B.H., and F.Y. performed the experiments. T.X. performed the modelling and computational work. X.C. and B.H. performed biochemical work. X.C. and F.Y. determined the cryo-EM structure of SLC44A1. All authors analyzed the data. T.X. and X.C. wrote the original draft with critical input from all authors. J.H. and Q.Z. edited the manuscript. J.H., Q.Z., T.X., and X.C. reviewed the final version.

## Competing interests

The authors declare no competing interests.

## Data and materials availability

All input data are available from public sources. Sequences of the human SLC superfamily were downloaded from UniProt database, IDs of which are available in SLC TABLES (http://slc.bioparadigms.org/). For prediction methods, the input databases include UniRef90 v2019_10 (https://ftp.ebi.ac.uk/pub/databases/uniprot/previous_releases/release-2019_10/uniref/), UniRef30_2020_03 (http://wwwuser.gwdg.de/~compbiol/uniclust/2020_03/), Uniclust30 v2018_08 (https://wwwuser.gwdg.de/~compbiol/uniclust/2018_08/), metagenome data metaclust_50 in Nov. 2020 (https://metaclust.mmseqs.org/current_release/). Atomic coordinates and EM density maps of SLC44A1 have been deposited in the Protein Data Bank (http://www.rcsb.org) and the Electron Microscopy Data Bank (https://www.ebi.ac.uk/pdbe/emdb/). Source code for the computation part of the protocol and the complete SLC ranking results are available under an open-source license at https://github.com/JingHuangLab/SLCFold. Correspondence and requests for materials should be addressed to (J.H.) and (Q.Z.).

## Notes

### Competing Interest Statement

The authors have declared no competing interest.

### Summary of Updates

Results updated with the cryo-EM structure of SLC44A1 obtained at a resolution of 3.86 A.

## Reference

1 Hediger, M. A. et al. The ABCs of solute carriers: physiological, pathological and therapeutic implications of human membrane transport proteinsIntroduction. Pflugers Arch 447, 465–468, doi:10.1007/s00424-003-1192-y (2004).

2 Hediger, M. A., Clémençon, B., Burrier, R. E. & Bruford, E. A. The ABCs of membrane transporters in health and disease (SLC series): introduction. Mol Aspects Med 34, 95–107, doi:10.1016/j.mam.2012.12.009 (2013).

3 Portugaly, E. & Linial, M. Estimating the probability for a protein to have a new fold: A statistical computational model. P Natl Acad Sci USA 97, 5161–5166, doi:10.1073/pnas.090559497 (2000).

4 Brenner, S. E. Target selection for structural genomics. Nat Struct Biol 7, 967–969, doi:10.1038/80747 (2000).

5 Portugaly, E., Kifer, I. & Linial, M. Selecting targets for structural determination by navigating in a graph of protein families. Bioinformatics 18, 899–907, doi:10.1093/bioinformatics/18.7.899 (2002).

6 Sasson, O. & Linial, M. ProTarget: automatic prediction of protein structure novelty. Nucleic Acids Res 33, W81–W84, doi:10.1093/nar/gki389 (2005).

7 McGuffin, L. J. & Jones, D. T. Targeting novel folds for structural genomics. Proteins 48, 44–52, doi:10.1002/prot.10129 (2002).

8 Wang, S., Sun, S. Q., Li, Z., Zhang, R. Y. & Xu, J. B. Accurate De Novo Prediction of Protein Contact Map by Ultra-Deep Learning Model. Plos Comput Biol 13, doi:10.1371/journal.pcbi.1005324 (2017).

9 Jumper, J. et al. Highly accurate protein structure prediction with AlphaFold. Nature, doi:10.1038/s41586-021-03819-2 (2021).

10 Senior, A. W. et al. Improved protein structure prediction using potentials from deep learning. Nature 577, 706–710, doi:10.1038/s41586-019-1923-7 (2020).

11 Baek, M. et al. Accurate prediction of protein structures and interactions using a three-track neural network. Science, eabj8754, doi:10.1126/science.abj8754 (2021).

12 Tunyasuvunakool, K. et al. Highly accurate protein structure prediction for the human proteome. Nature, doi:10.1038/s41586-021-03828-1 (2021).

13 Burley, S. K. et al. Protein Data Bank: the single global archive for 3D macromolecular structure data. Nucleic Acids Res 47, D520–D528, doi:10.1093/nar/gky949 (2019).

14 Yang, J. Y. et al. Improved protein structure prediction using predicted interresidue orientations. P Natl Acad Sci USA 117, 1496–1503, doi:10.1073/pnas.1914677117 (2020).

15 Xu, J., Mcpartlon, M. & Li, J. Improved protein structure prediction by deep learning irrespective of co-evolution information. bioRxiv, 2020.2010.2012.336859, doi:10.1101/2020.10.12.336859 (2020).

16 Bai, X. Y., Moraes, T. F. & Reithmeier, R. A. F. Structural biology of solute carrier (SLC) membrane transport proteins (2018). Mol Membr Biol 34, 65–65, doi:10.1080/09687688.2018.1503851 (2017).

17 Perland, E. & Fredriksson, R. Classification Systems of Secondary Active Transporters. Trends Pharmacol Sci 38, 305–315, doi:10.1016/j.tips.2016.11.008 (2017).

18 Bai, X. Progress in Structural Biology of Solute Carriers. Current Molecular Biology Reports 7, 9–19, doi:10.1007/s40610-021-00144-5 (2021).

19 Holm, L. Benchmarking fold detection by DaliLite v.5. Bioinformatics 35, 5326–5327, doi:10.1093/bioinformatics/btz536 (2019).

20 Luscombe, N. M., Qian, J. A., Zhang, Z. L., Johnson, T. & Gerstein, M. The dominance of the population by a selected few: power-law behaviour applies to a wide variety of genomic properties. Genome Biol 3, doi:10.1186/gb-2002-3-8-research0040 (2002).

21 Tsai, J. Y. et al. Structure of the sodium-dependent phosphate transporter reveals insights into human solute carrier SLC20. Sci Adv 6, doi:10.1126/sciadv.abb4024 (2020).

22 Zhang, Y. & Skolnick, J. TM-align: a protein structure alignment algorithm based on the TM-score. Nucleic Acids Res 33, 2302–2309, doi:10.1093/nar/gki524 (2005).

23 Assur Sanghai, Z. et al. Structure-based analysis of CysZ-mediated cellular uptake of sulfate. Elife 7, doi:10.7554/eLife.27829 (2018).

24 Bernsel, A., Viklund, H., Hennerdal, A. & Elofsson, A. TOPCONS: consensus prediction of membrane protein topology. Nucleic Acids Res 37, W465–W468, doi:10.1093/nar/gkp363 (2009).

25 Xu, J., McPartlon, M. & Li, J. Improved protein structure prediction by deep learning irrespective of co-evolution information. Nat Mach Intell 3, 601–609, doi:10.1038/s42256-021-00348-5 (2021).

26 Li, Y. et al. Deducing high-accuracy protein contact-maps from a triplet of coevolutionary matrices through deep residual convolutional networks. Plos Comput Biol 17, e1008865, doi:10.1371/journal.pcbi.1008865 (2021).

27 Wang, S., Peng, J., Ma, J. Z. & Xu, J. B. Protein Secondary Structure Prediction Using Deep Convolutional Neural Fields. Sci Rep-Uk 6, doi:10.1038/srep18962 (2016).

28 Kabsch, W. & Sander, C. Dictionary of protein secondary structure: pattern recognition of hydrogen-bonded and geometrical features. Biopolymers 22, 2577–2637, doi:10.1002/bip.360221211 (1983).

29 Yamashita, A., Singh, S. K., Kawate, T., Jin, Y. & Gouaux, E. Crystal structure of a bacterial homologue of Na+/Cl--dependent neurotransmitter transporters. Nature 437, 215–223, doi:10.1038/nature03978 (2005).

30 Lei, J. & Frank, J. Automated acquisition of cryo-electron micrographs for single particle reconstruction on an FEI Tecnai electron microscope. J Struct Biol 150, 69–80, doi:10.1016/j.jsb.2005.01.002 (2005).

31 Zheng, S. Q. et al. MotionCor2: anisotropic correction of beam-induced motion for improved cryo-electron microscopy. Nat Methods 14, 331–332, doi:10.1038/nmeth.4193 (2017).

32 Grant, T. & Grigorieff, N. Measuring the optimal exposure for single particle cryo-EM using a 2.6 A reconstruction of rotavirus VP6. Elife 4, e06980, doi:10.7554/eLife.06980 (2015).

33 Zhang, K. Gctf: Real-time CTF determination and correction. J Struct Biol 193, 1–12, doi:10.1016/j.jsb.2015.11.003 (2016).

34 Zivanov, J. et al. New tools for automated high-resolution cryo-EM structure determination in RELION-3. Elife 7, doi:10.7554/eLife.42166 (2018).

35 Wang, N. et al. Structural basis of human monocarboxylate transporter 1 inhibition by anti-cancer drug candidates. Cell 184, 370–383 e313, doi:10.1016/j.cell.2020.11.043 (2021).

36 Rosenthal, P. B. & Henderson, R. Optimal determination of particle orientation, absolute hand, and contrast loss in single-particle electron cryomicroscopy. J Mol Biol 333, 721–745 (2003).

37 Scheres, S. H. & Chen, S. in Nature methods Vol. 9 853–854 (2012).

38 Chen, S. et al. High-resolution noise substitution to measure overfitting and validate resolution in 3D structure determination by single particle electron cryomicroscopy. Ultramicroscopy 135, 24–35, doi:10.1016/j.ultramic.2013.06.004 (2013).

39 Trabuco, L. G., Villa, E., Mitra, K., Frank, J. & Schulten, K. Flexible fitting of atomic structures into electron microscopy maps using molecular dynamics. Structure 16, 673–683, doi:10.1016/j.str.2008.03.005 (2008).

40 Adams, P. D. et al. PHENIX: a comprehensive Python-based system for macromolecular structure solution. Acta Crystallogr D Biol Crystallogr 66, 213–221, doi:10.1107/S0907444909052925 (2010).

41 Emsley, P., Lohkamp, B., Scott, W. G. & Cowtan, K. Features and development of Coot. Acta Crystallogr D Biol Crystallogr 66, 486–501, doi:10.1107/S0907444910007493 (2010).

