## Supplemental Information for "Rational Exploration of Fold Atlas for Human Solute Carrier Proteins"

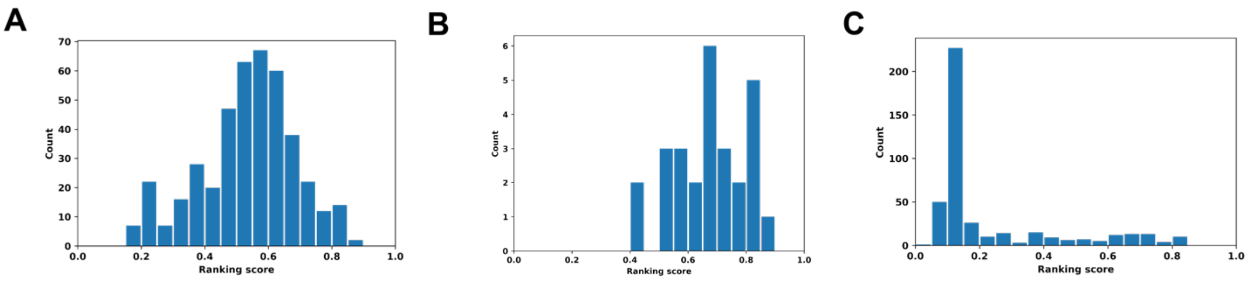


**Fig. S1. Validation of the fitness score.** Histogram of the fitness ranking scores for all of 425 SLC members (**A**) and structural-known SLC members (**B**) relative to all known membrane proteins. (**C**) Distribution of the fitness ranking scores for 425 target proteins assuming only LeuT fold structures are known.

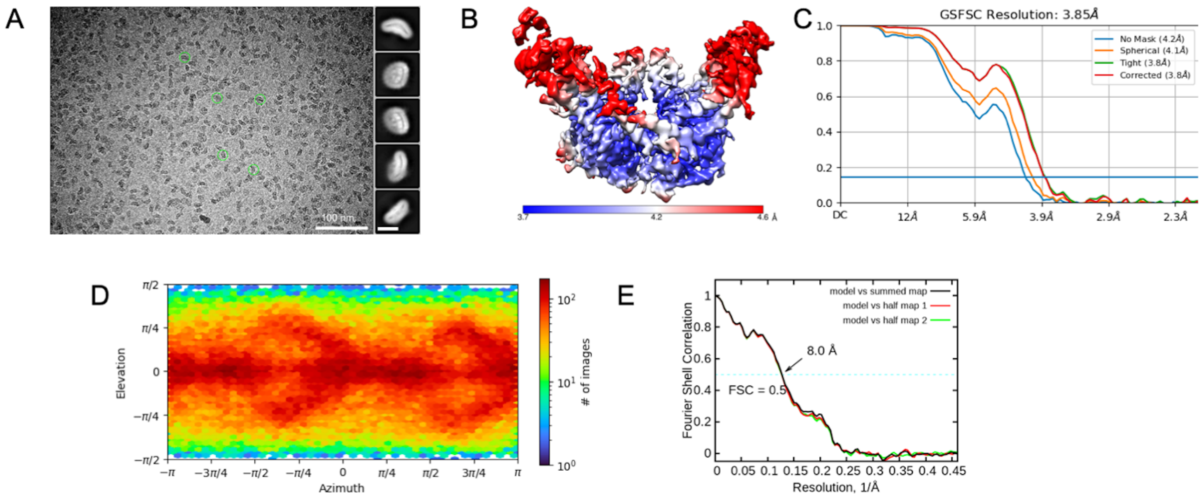


**Fig. S2. Cryo-EM analysis of SLC44A1.** (**A**) Representative cryo-EM micrographs and 2D average results. (**B**) Local resolution map for the 3D reconstruction. (**C**) Gold standard FSC curves for the 3D refinements applied with protomer mask. (**D**) Euler angle distribution for the 3D reconstruction. (**E**) FSC curve of the refined model of SLC44A1 versus the overall map that it is refined against; FSC curve of the model refined against the first half map versus the same map (red); and FSC curve of the model refined against the first half map versus the second half map (green). The small difference between the red and green curves indicates that the refinement of the atomic coordinates did not suffer from overfitting.

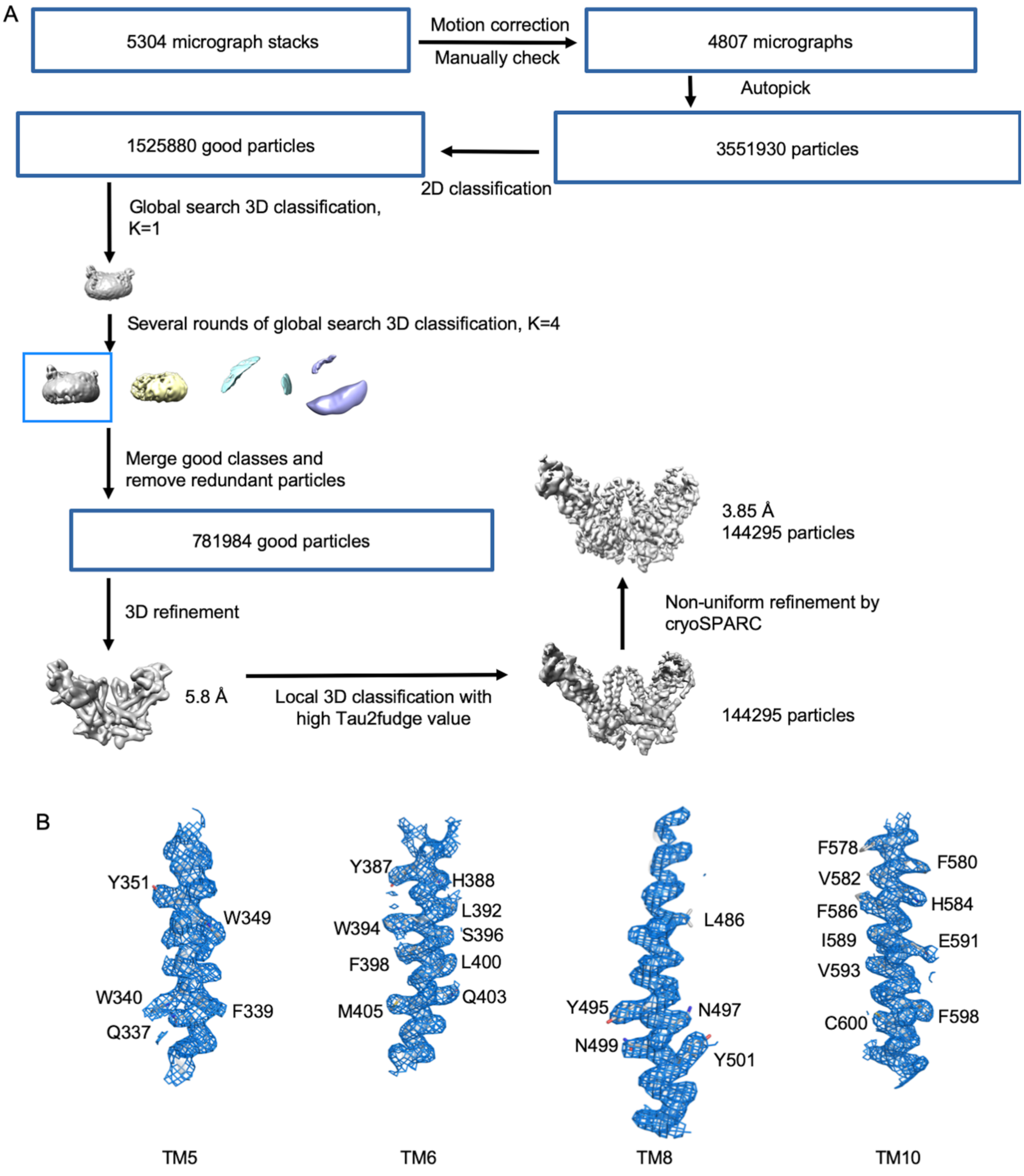


**Fig. S3. Flowchart for cryo-EM data processing and representative regional maps**. (**A**) Flowchart for cryo-EM data processing, please refer to the 'Cryo-EM sample preparation, data acquisition and procession' section in Methods for details. (**B**) electron-density map of the indicated transmembrane (TM) regions. The quality of the map is good enough for identification of all the transmembrane helices.

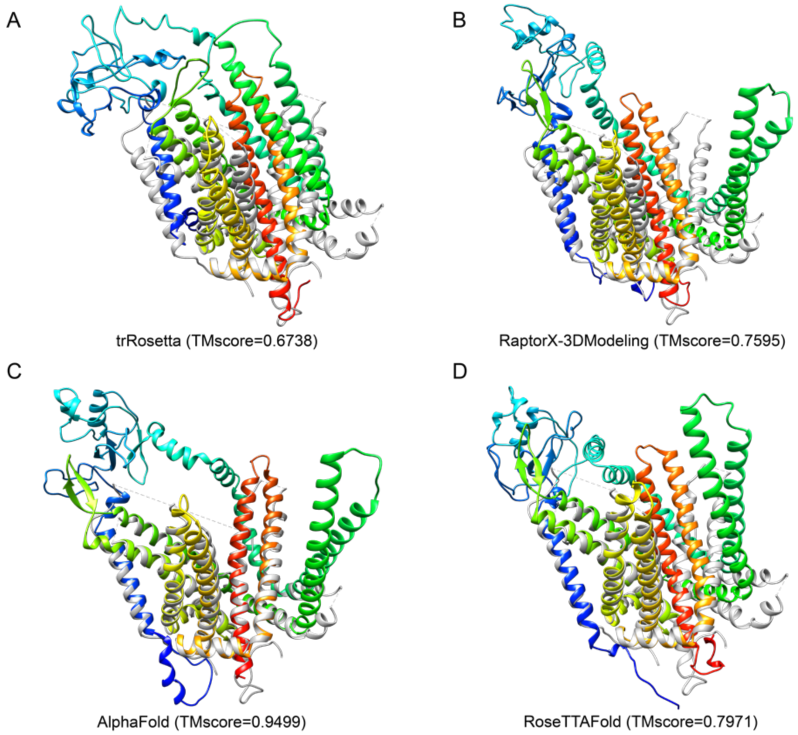


**Fig. S4. Comparisons of predicted structures (rainbow) to the build model from cryo-EM map (grey).**

**Table S1.** The fitness score and most similar structures of 15 SLC proteins ranked with highest *S* scores.

| SLC gene | *S* | PDB ID of the most similar structure | Gene name | Homologs/SLC family/Gene |
|---|---|---|---|---|
| SLC6A8 | 0.8033 | 6VRL | SLC6A4 | SLC6 |
| SLC26A9 | 0.8056 | 6RTF | Slc26A9 | Homologs |
| SLC6A6 | 0.8154 | 6DZY | SLC6A4 | SLC6 |
| SLC6A7 | 0.8185 | 6VRL | SLC6A4 | SLC6 |
| SLC2A5 | 0.8193 | 7CRZ | SLC2A3 | SLC2 |
| SLC2A11 | 0.8231 | 7CRZ | SLC2A3 | SLC2 |
| SLC6A4 | 0.8278 | 5I75 | SLC6A4 | SLC6A4 |
| SLC6A1 | 0.8325 | 6VRL | SLC6A4 | SLC6 |
| SLC6A3 | 0.8388 | 5I75 | SLC6A4 | SLC6 |
| SLC2A1 | 0.8399 | 7CRZ | SLC2A3 | SLC2 |
| SLC6A11 | 0.8415 | 6DZY | SLC6A4 | SLC6 |
| SLC6A13 | 0.8455 | 6VRL | SLC6A4 | SLC6 |
| SLC14A1_UT-B1 | 0.8498 | 6QD5 | SLC14A1 | SLC14A1 |
| SLC2A3 | 0.8521 | 7CRZ | SLC2A3 | SLC2A3 |
| SLC6A12 | 0.8545 | 6DZY | SLC6A4 | SLC6 |

"PDB ID of the most similar structure" is the ID of the structure most similar with the 3D structure predicted by trRosetta among all known structures in RCSB PDB. "Gene name" refers to gene name of those PDB structures. "Homologs/SLC family/Gene" refers to the common ground between a SLC member and the most similar structure, where "Homologs" means the two proteins are homologous, "SLC family" means the two proteins belong to a SLC family but are different genes, and "Gene" means the two structures correspond to the same gene.

**Table S1 | SLC Fold information in this work.** The IDs of the newly recognized SLC members are shown in orange and bold.

| Fold names | No. TMs | Fold features | PDB ID of Rep. | No. families | No. SLCs | Year | PDB ID/chain ID of the earliest structure | SLC family IDs | Fold name previously |
|---|---|---|---|---|---|---|---|---|---|
| MFS | 12 | TMs 1-6 and TMs 7-12 are "6+6" inverted repeats; TMs 1-3 and TMs 4-6 are "3+3" inverted repeats; TM 4 is surround by TMs 1-3; TMs 1, 2, 4, 5 form the repeat interface while TMs 3, 6 locates in the distance. TMs 1-3 are arranged in a left-handed spiral mode. | 1PV6 | 21 | 119 | 2003 | 1PV6/A | 2, 15-19, 22, 29, 33, 37, 40, 43, 45, 46, 49, **52**, **59**, **60**, **61**, 63, O | MFS |
| LeuT | 10 | TMs 1-5 and TMs 6-10 are "5+5" inverted repeats; there are unwound regions in the middle of TM 1 and TM 6; between TMs 2-3 and TMs 7-8 locate soluble short helices. Other TMs are in the peripheral area. | 6TL2 | 8 | 71 | 2008 | 3DH4/A | 5-7, 11, 12, 32, 36, 38 | LeuT |
| Aac | 6 | TMs (1, 6), TMs (2,3), and TMs (4, 5) are "2+2+2" rotated repeats. | 4C9J | 1 | 53 | 2014 | 4C9G/A | 25 | NA |
| YddG | 10 | TMs 1-5 and TMs 6-10 are "5+5" inverted repeats; TMs 1-4 and TMs 6-9 form central cavity, beside which locate TM 5 and TM 10. | 6UKJ | 1 | 30 | 2016 | 5I20/A | 35 | NA |

| | | | | | | | | | |
|---|---|---|---|---|---|---|---|---|---|
| UraA | 14 | TMs 1-7 and TMs 8-10 are "7+7" inverted repeats; there are unwound regions in the middle of TMs 3 and 10; TMs 1-4 and TMs 8-11 form core domain; TMs 5-7 and TMs 12-14 are the scaffold domain. | 5I6C | 3 | 24 | 2011 | 3QE7/A | 4, 23, 26 | NA |
| NhaA | 10 | TMs 1-5 and TMs 6-10 are inverted repeats; TM4 and TM8 are partially unwound in the middle. TMs 3-5 and TMs 8-10 are the core domain; TMs 1, 2, 6, and 7 are the scaffold domain; the interface of the two domains is composed of TMs 3, 4, 8, and 9. | 7CYK | 2 | 20 | 2009 | 3FI1/A | 9, 10 | CPA/AT |
| ZIP | 8 | TMs 1-3 and TMs 6-8 are inverted repeats. TMs 4, 5 are sandwiched by the two 3-TM repeats. | 6PGI | 1 | 14 | 2017 | 5TSB/A | **39** | NA |

| | | | | | | | | | |
|---|---|---|---|---|---|---|---|---|---|
| CNT | 10 | TMs 1-5 and TMs 6-10 are "5+5" inverted repeats; the first three TMs from N terminal are marked as -2, -1, 0; TMs 1 and 6 are two long, slanted, and bent helices; TMs 2-3 and TMs 7-8 form hairpin; there are unwound regions in the middle of TMs 4 and 9; TMs 3, 4, the end of TM 2, and the start of TM 5 form right-handed spirals. | 3TIJ | 2 | 11 | 2012 | 3TIJ/A | 13, **28**, **34** | NA |
| YiiP | 6 | It functions as a dimer. TMs 1-3 and TMs 4-6 are "3+3" rotated repeats and face with each other. | 7KZZ | 1 | 10 | 2009 | 3H90/A | 30 | NA |
| NCX | 10 | TMs 1-5 and TMs 6-10 are inverted "5+5" repeats; TMs 3-5 and TMs 8-10 are core domain; TMs 1, 2, 6, and 7 are the scaffold domain. | 3V5U | 2 | 9 | 2012 | 3V5U/A | 8, **24** | CaCA |
| GltpH | 12 | Two "3+3" inverted repeats: TMs 1-3 and TMs 4-6; TMs 7-9 and TMs 10-12; TMs 1, 2, 4, 5 are core domain; TMs 1-5 are the scaffolding; TMs 7-8 and TMs 10-11 forms two hairpins; unwound or twisted regions exist in the middle of TMs 9 and 12. | 6WYL | 1 | 7 | 2007 | 2NWL/A | 1 | Glt |

| | | | | | | | | | |
|---|---|---|---|---|---|---|---|---|---|
| NorM | 12 | TMs 1-6 and TMs 7-12 are "6+6" rotated repeats. Each repeat forms a bundle domain. TMs 1, 2, 7, and 8 are the domain interface. TMs 3, 4, 5, and 6 locates along a line that surrounds TM1-2. TM 6 is inserted in the between of TMs 4 and 5. | 5Y50 | 3 | 6 | 2010 | 3MKU/A | **41**, 47, **62** | MviN, MATE |
| AmtB | 10 | The first and last TMs are marked as 0 and 11. TMs 1-5 and TMs 6-10 are "5+5" inverted repeats. Each repeat form left-handed spirals. TMs 1, 3, 5, 6, 8, and 10 locates at the inter-domain interface. TMs 1 and 11 are locate at the beginning and the end of the fold. | 3HD6 | 2 | 5 | 2004 | 1U77/A | 14, 42 | Channel like |
| SLC44 | 10 | TMs 1 and 2 are on the outside of the protein. TMs 3-10 are core domain with 4 groups of two joint TMs. Both TMs (6, 7) and TMs (8, 9) are linked by a helix. | NA | 1 | 5 | 2021 | NA | **44** | NA |

| | | | | | | | | | |
|---|---|---|---|---|---|---|---|---|---|
| AcrB | 12 | The first TM is marked as 0. TMs 1-6 and TMs 7-12 are “6+6” rotated repeats. TMs 1 and 7 both have a right-angle bend, half of which is along with the membrane surface. TMs 2, 4, and 5 are the interface. TMs 2-6 forms resistance nodulation division fold. | 6V3H | 1 | 2 | 2005 | 1T9W/A | **65** | NA |
| PiT | 12 | TMs 1-5 and TMs 8-12 are “5+5” inverted repeats. TMs 6 and 7 are inserted in the two repeats. TMs 3-4 and TM 8-9 are two hairpins, where substrate bound. | 6L85 | 1 | 2 | 2020 | 6L85/A | **20** | NA |
| SWEET 2b | 7 | TMs 1-3 and TMs 5-7 are rotated repeats, linked by TM 4. | 5XPD | 1 | 1 | 2015 | 5TCT/A | 50 | MtN3-like |
| bacteriorhodopsin | 7 | TMs 1-4 and TM5-7 are located in different planes, and both behave zigzag shape. | 5W0P | 1 | 1 | 1999 | 1C3W/A | **51(A)** | NA |
| SLC64 | 6 | TMs 1-3 and TMs 4-6 are inverted repeats linked by a long helix, which is parallel to the membrane plane. | NA | 1 | 1 | NA | NA | **64** | NA |
| SLC57 | 10 | TMs 1-5 and TMs 6-10 are inverted repeats. TM 3 is inserted in the between of TM 1 and TM 2. | NA | 1 | 6 | NA | NA | **57** | NA |

| | | | | | | | | | |
|---|---|---|---|---|---|---|---|---|---|
| SLC53 | 10 | 10 TMs without structural repeats. | Part of 6QQ6 | 1 | 1 | NA | NA | **53** | NA |
| SLC31 | 9 | Trimer of SLC31 by GalaxyHomomer server[1] is similar with Ctr1(PDB ID: 6M97)[2]. | 6M97 | 1 | 2 | NA | NA | **31** | NA |
| SLC56 | 6 | TMs 1-2, TMs 2-3, and TMs 4-5 are "2+2+2" repeats. It is a channel-like fold. | 6C70 | 1 | 5 | NA | Part of 1T9W | **56** | NA |
| 5TMs | 5 | SLC58 members have a partial experimental structure (PDB ID: 6S7T) and functions as heteromer. | NA | 1 | 2 | NA | NA | 58 | NA |
| 4TMs | 4 | 4 TM helices | NA | 2 | 2 | NA | NA | 35(E2), 48 | NA |
| 3TMs | 3 | 3 TM helices | NA | 1 | 6 | NA | NA | 27 | NA |
| 1TM | 1 | SLC51B, SLC54, SLC55 function in complex. SLC3 members are chaperons. | NA | 4 | 9 | NA | NA | 3, 51(B), 54, 55 | NA |

"Fold names" are the gene names of the protein belonging to the fold, which was first experimentally resolved, or the number of TMs when the number of TM helices is less than 6, or SLC family ID. "PDB ID of Rep." refers to the PDB ID of the representative structure for a SLC member. "Year" refers to the year that a fold was resolved at the first time.

**Table S3.** Ranking score of the top 35 SLC members that probably have a new fold.

| No. | SLC | $S_{3D}$ | $C_{3D}$ | $\overline{C_{2D}}$ | $\overline{C_{1D}}$ | $S$ |
|---|---|---|---|---|---|---|
| 1 | SLC44A5 | 0.3738 | 0.6284 | 0.5272 | 0.7818 | 0.1557 |
| 2 | SLC44A4 | 0.3844 | 0.6435 | 0.5401 | 0.8050 | 0.1630 |
| 3 | SLC39A4 | 0.3907 | 0.5317 | 0.4096 | 0.8889 | 0.1755 |
| 4 | SLC44A2 | 0.4040 | 0.6251 | 0.5474 | 0.8072 | 0.1807 |
| 5 | SLC44A1 | 0.4169 | 0.6463 | 0.5739 | 0.8432 | 0.1896 |
| 6 | SLC55A2 | 0.3992 | 0.5414 | 0.2454 | 0.8204 | 0.1904 |
| 7 | SLC44A3 | 0.4167 | 0.6168 | 0.5714 | 0.8162 | 0.1918 |
| 8 | SLC34A3 | 0.4369 | 0.7675 | 0.5289 | 0.8103 | 0.2029 |
| 9 | SLC34A2 | 0.4295 | 0.6570 | 0.5078 | 0.7755 | 0.2044 |
| 10 | SLC39A10 | 0.3208 | 0.2777 | 0.2389 | 0.8676 | 0.2092 |
| 11 | SLC8A3 | 0.3572 | 0.4359 | 0.5680 | 0.7708 | 0.2095 |
| 12 | SLC55A3 | 0.4396 | 0.7086 | 0.3194 | 0.8140 | 0.2142 |
| 13 | SLC64A1 | 0.4428 | 0.6548 | 0.5448 | 0.8469 | 0.2142 |
| 14 | SLC58A2 | 0.4349 | 0.5030 | 0.5977 | 0.8040 | 0.2149 |
| 15 | SLC34A1 | 0.4471 | 0.6861 | 0.5432 | 0.8078 | 0.2176 |
| 16 | SLC8A1 | 0.3673 | 0.3988 | 0.5627 | 0.7739 | 0.2177 |
| 17 | SLC25A13 | 0.3641 | 0.4278 | 0.4348 | 0.8248 | 0.2178 |
| 18 | SLC39A7 | 0.4413 | 0.5943 | 0.4648 | 0.8795 | 0.2180 |
| 19 | SLC24A3 | 0.4388 | 0.5388 | 0.5303 | 0.8131 | 0.2191 |
| 20 | SLC24A4 | 0.4457 | 0.6166 | 0.5643 | 0.8262 | 0.2194 |
| 21 | SLC39A6 | 0.3435 | 0.3397 | 0.2411 | 0.8532 | 0.2199 |
| 22 | SLC24A2 | 0.4419 | 0.5551 | 0.5248 | 0.8343 | 0.2205 |
| 23 | SLC8A2 | 0.3848 | 0.4500 | 0.6068 | 0.7932 | 0.2217 |
| 24 | SLC25A12 | 0.3882 | 0.4886 | 0.4196 | 0.8167 | 0.1777 |
| 25 | SLC24A1 | 0.3149 | 0.1043 | 0.1215 | 0.7953 | 0.2303 |
| 26 | SLC53A1 | 0.4586 | 0.6671 | 0.4690 | 0.8203 | 0.2320 |
| 27 | SLC39A12 | 0.3872 | 0.4305 | 0.3531 | 0.8405 | 0.2348 |
| 28 | SLC14A2_UT-A1 | 0.4052 | 0.4464 | 0.5180 | 0.8194 | 0.2370 |
| 29 | SLC25A24 | 0.4695 | 0.6115 | 0.5587 | 0.8497 | 0.2431 |
| 30 | SLC58A1 | 0.4417 | 0.4925 | 0.5940 | 0.8087 | 0.2516 |

### Reference


1 Baek, M., Park, T., Heo, L., Park, C. & Seok, C. GalaxyHomomer: a web server for protein homo-oligomer structure prediction from a monomer sequence or structure. *Nucleic Acids Res* **45**, W320-W324, doi:10.1093/nar/gkx246 (2017).

2 Ren, F. *et al.* X-ray structures of the high-affinity copper transporter Ctr1. *Nat Commun* **10**, doi:10.1038/s41467-019-09376-7 (2019).
